# The expression of integron arrays is shaped by the translation rate of cassettes

**DOI:** 10.1101/2024.03.26.586746

**Authors:** André Carvalho, Alberto Hipólito, Filipa Trigo da Roza, Lucía García-Pastor, Ester Vergara, Aranzazu Buendía, Teresa García-Seco, José Antonio Escudero

## Abstract

Integrons are key elements in the rise and spread of multidrug resistance in Gram-negative bacteria. These genetic platforms capture cassettes containing promoterless genes and stockpile them in arrays of variable length. In the current integron model, expression of cassettes is granted by the Pc promoter in the platform and is assumed to decrease as a function of its distance. Here we explored this model using a large collection of 136 antibiotic resistance cassettes and show that the effect of distance is in fact negligible. Instead, cassettes have a strong impact in the expression of downstream genes because their translation rate affects the stability of the whole polycistronic mRNA molecule. Hence, poorly translated cassettes decrease the expression and resistance phenotype of cassettes downstream. Our data puts forward a novel integron model in which expression is contingent on the translation of cassettes upstream, rather than on the distance to the Pc.

## INTRODUCTION

Antimicrobial resistance (AMR) is one of the major threats for health globally^1^. The spread of resistance genes through Horizontal Gene Transfer (HGT) has fostered the rise of AMR during the last decades. Mobile Integrons (MIs) have been key players in this phenomenon through their association with mobile genetic elements^2–4^. MIs are genetic platforms that capture and stockpile new genes through site specific recombination. Indeed, 89% of cassettes in MIs encode antimicrobial resistance genes against a variety of antibiotic families (IntegrAll database,^5^).

Structurally, integrons are genetic elements composed of a stable platform and a variable array of genes embedded in discrete elements called integron cassettes. The stable platform comprises an integrase-coding gene (*intI*), as well as a recombination site (*attI*) for cassette integration. Cassettes are generally composed of an open reading frame (ORF) and an *attC* recombination site. Arrays in MIs can contain from one up to eleven cassettes. In class 1 integrons, the majority (80%) of arrays contain 1 to 3 cassettes (IntegrAll database,^5^). Importantly cassettes are generally promoterless, and their expression is fostered by the dedicated Pc promoter, located within the platform, upstream of the *attI* site where cassettes are inserted. This makes of integron arrays an operon-like structure, with a single promoter governing the expression of several cassettes. As a consequence, it is generally accepted that cassettes at first position in the array are more expressed than those further downstream, simply as a function of the distance to the promoter. Nevertheless, the order of cassettes is not static, since the integrase is able to re-shuffle them, excising and integrating them in the first position of the array. Because the integrase is under the control of the host’s SOS response ^6^, integrons represent a low-cost memory of functions that provides adaptation on demand ^7–10^ (Fig. 1A).

**Figure 1.**
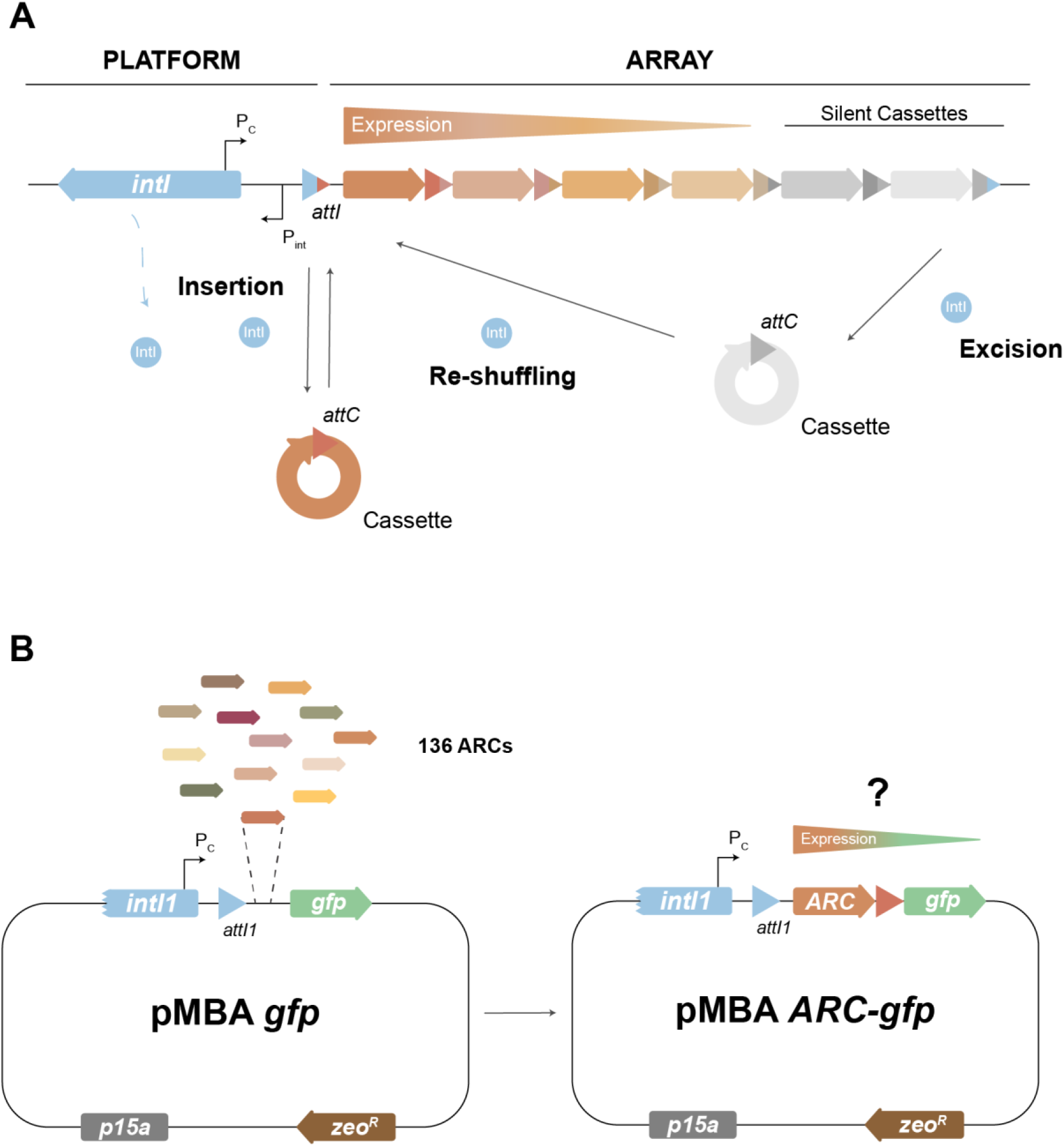
**(A)** Schematic representation of the current integron model. When expressed, the integrase (encoded by the *intI* gene) is able to integrate, excise and re-shuffle discrete elements called cassettes. These are composed of a gene and an *attC* recombination site, and are arranged in arrays that contain multiple cassettes. Cassettes are generally promoterless but can be expressed when integrated at the *attI* site, where their expression is controlled by the integron-borne Pc promoter. An expression gradient is then generated: the first cassette displays the highest expression while expression gradually decreases for cassettes further away in the array. **(B)** 136 different antibiotic resistance cassettes were individually cloned in first position in pMBA vector^20,21^, allowing to study how each ARC affects the expression gradient.

The effect of cassette position has been observed in several studies ^10–13^ so that the distance-to-Pc gradient of expression along the array is a paradigm that has been central to the working model of integrons. Only some exceptions to this rule have been documented, like the existence of cassettes that contain their own promoters ^14–18^ or the presence of small ORFs in *attC* sites that enhance translation of downstream cassettes ^13^. However, most of these examples only cover a few integron cassettes and seem to remain anecdotal. Recently, we have also shown the existence of gene-less cassettes in chromosomal integrons that contain promoters modulating the expression of the array ^19^. All these mechanisms suggest that expression in integrons does not always fit the distance-to-Pc gradient. This motivated us to revisit the expression model of the integron, and specifically, if and how cassettes shape the expression of the array and influence the resistance conferred by others <u>a</u>ntibiotic resistance <u>c</u>assettes (ARCs).

In this work, we study the effect of 136 ARCs on the expression of a downstream *gfp* gene. We show that the impact of a cassette in downstream expression is independent of and more important than the distance to the Pc. This impact is strong enough to decrease resistance levels conferred by a second ARC below clinical breakpoints. To determine the mechanism underlying this phenomenon, we first assessed the influence of known exceptions to the rule of distance, and found an occasional and generally modest contribution of additional promoters or *attC* sites. Instead, the translation rate of the first ARC influences the expression of the second cassette by altering the mRNA levels. This effect is strong and is pervasive across the collection. Our findings change the working model of the integron, to one where the expression gradient of the array becomes dependent on the identity of the first cassette.

## RESULTS

### Cassette identity modulates expression of the array downstream

To assess if cassettes can modulate the expression of downstream genes in the array (i.e.: if they have polar effects on the array) we took advantage of the recently generated pMBA collection ^20,21^. pMBA is a p15a replicon designed to provide cassettes with an appropriate genetic context: it contains a class 1 integron platform encoding the Pc, the Pint, and the *attI site* where cassettes are cloned in first position mimicking an integrase mediated *attI* x *attC* reaction. The integrase gene (*intI1*) is truncated to avoid its well-known deleterious effect ^22^. Immediately downstream each cassette there is a *gfp* gene mimicking a second cassette (Fig. 1B) ^20^. The collection is composed of 136 pMBA variants containing different ARCs in *E. coli* MG1655. Transcription is driven by the strong variant of the Pc promoter (PcS). Hence, the pMBA collection offers an isogenic setting to study the impact of 136 cassettes on the rest of the array (Fig. 1B). Hereafter, and for the sake of simplicity, we will use the name of the resistance gene encoded in each cassette to refer to the corresponding *E. coli*-pMBA-ARC variant.

We measured GFP fluorescence in all 136 *E. coli* strains in the collection using flow cytometry. Data was then normalized to the fluorescence of the *E. coli* strain containing pMBAø, which carries the *gfp* gene in the first position in the array. Our results show a large variation in the expression of the second cassette across the collection (Fig. 2A), with fluorescence levels scattered across the 120-fold range between the most and the least repressive ones (*bla*_OXA-20_ and *aacA43* respectively). A similarly broad distribution of fluorescence values can be observed for each different antibiotic family represented in the collection, ruling out the influence of the function encoded (Supplementary figure 1). As expected, the vast majority of ARCs decrease the expression of the second cassette, with a mean fluorescence ratio (pMBA-ARC / pMBAø) of 0.417 for all ARCs (Fig. 2B). While this value can be interpreted as the average repressive effect of cassettes on downstream genes, our data shows a broad dispersion, with some cassettes even increasing fluorescence mildly (up to 2-fold) (n=6, ≍4%).

**Figure 2.**
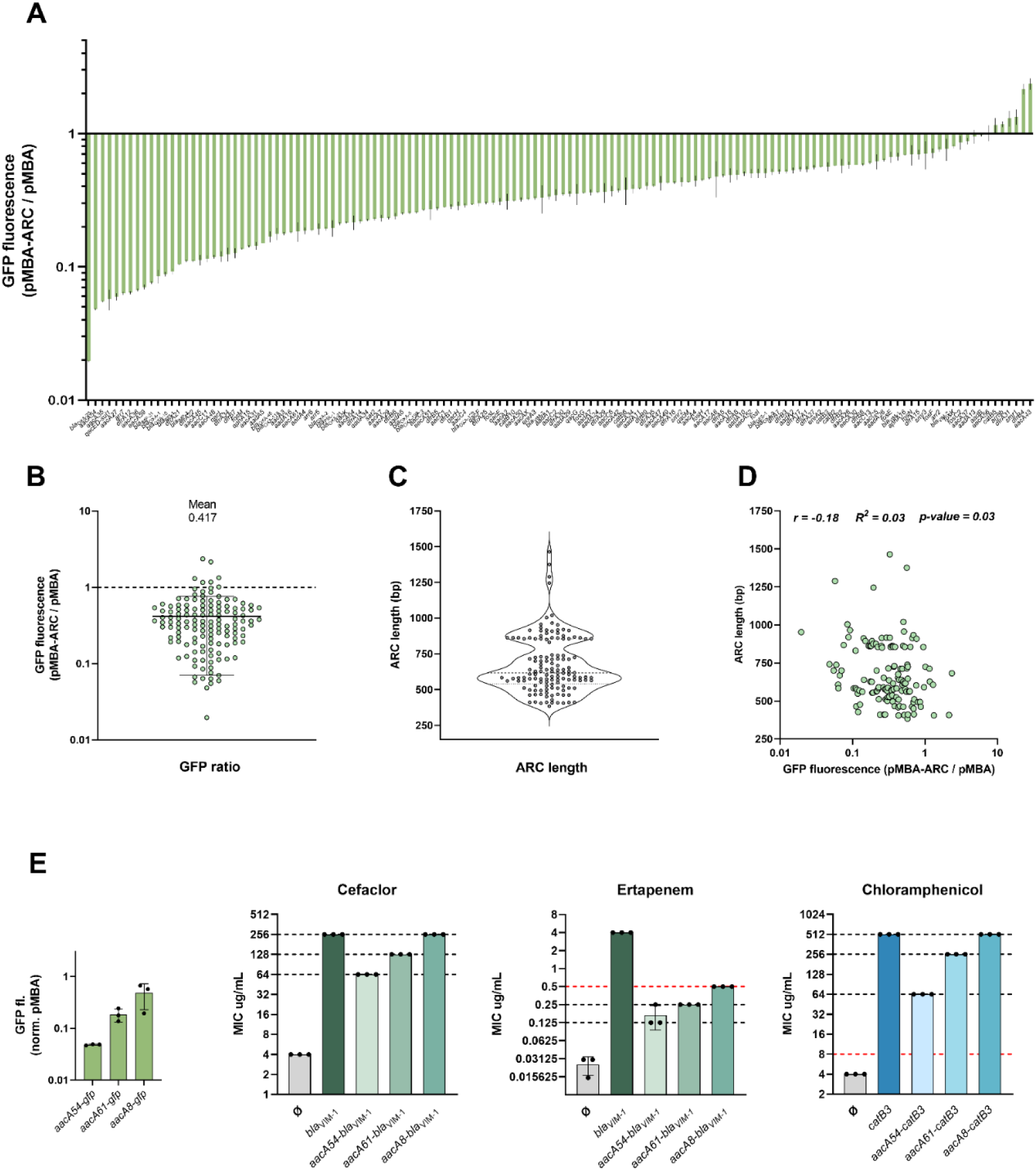
Cassette identity dictates expression and the resistance level of downstream ARCs. **(A)** GFP fluorescence of all 136 pMBA-ARC strains measured by flow cytometry normalized to fluorescence of pMBA control strain. Bars represent the mean and SD from three biological replicates. **(B)** Distribution of the GFP fluorescence ratios (pMBA-ARC / pMBA) with indication of the mean value and SD. **(C)** Violin plot depicting cassette length distribution of all 136 ARCs. Median and quartiles are represented. **(D)** Correlation between relative GFP fluorescence levels and ARC length. *r* indicates Pearson’s correlation coefficient. **(E)** Relative fluorescence of *aacA54*, *aacA61* and *aacA8* cassettes. Resistance levels conferred by pMBAØ or pMBA carrying the indicated arrays to cefaclor, ertapenem and chloramphenicol. Bars depict the mean and SD of the MIC values of three independent biological replicates. A red dotted line indicates the clinical breakpoint (EUCAST) for *E. coli* against the respective antibiotic.

Our data highlights that cassettes can exert very different -even opposite-effects on the expression of the array. This puts the model of expression of integrons in question. Our experimental setup allows for the verification of the distance-to-Pc model since the distance to the second cassette (the *gfp* gene) is the length of the ARC in first position. ARC length varies substantially in the pMBA collection, with the lower and upper limits being 384bp-(*dfrB2*) and 1464bp-long (*ereA3*) (Fig. 2C). However, correlation between ARC length and GFP fluorescence ratios was inexistent (r= -0.18, *p =* 0.03) (Fig. 2D). We conclude that the effect of the distance to the Pc is probably masked by other mechanisms, to the point of becoming negligible. Instead, our data shows that the expression of the second cassette depends on the identity of the first cassette.

To study how this could impact the resistance levels of downstream ARCs, we specifically selected three aminoglycoside resistance cassettes (*aacA54*, *aacA61*, and *aacA8*) displaying distinct levels of polar effects (Fig. 2E). We substituted the *gfp* gene in these strains with *bla*_VIM-1_ cassette and examined their resistance to β-lactams ertapenem and cefaclor (*aacA54*, *aacA61* or *aacA8* do not show cross resistance to these antibiotics (Supplementary figure 2)). Notably, *aacA54* and *aacA61* significantly reduced cefaclor resistance, while *aacA8* had no effect (Fig. 2E). All three cassettes strongly reduced ertapenem resistance, with the extent of reduction depending on cassette identity. Of note, *aacA54* and *aacA61* (but not *aacA8*) lowered ertapenem resistance below the clinical breakpoint, highlighting the clinical relevance of cassette identity. Moreover, this pattern persisted when we replaced the *bla*_VIM-1_ cassette with *catB3* and measured chloramphenicol resistance. Hence, polar effects in integrons arrays are contingent on cassette identity and are clinically relevant.

### Different ARCs modulate downstream array expression mostly by impacting mRNA levels

We next sought to investigate the underlying mechanisms modulating cassette expression. An ARC can affect the expression of the second cassette by changing its mRNA and/or protein levels. To assess to what extent each ARC impacts *gfp* mRNA levels, we performed qRT-PCR of the *gfp* gene in all 136 pMBA-ARC strains and calculated its fold change relative to pMBAø (Fig. 3A). Our results indicate that most ARCs affect negatively the mRNA levels of the *gfp* gene (mean mRNA fold change = 0.757) and that variation is similar to that observed for GFP fluorescence (Fig. 3B). mRNA levels showed a strong positive correlation (r = 0.73, R^2^ *= 0.53, p* < 0.001) with fluorescence ratios (Fig. 3C). It is of note that the values of *gfp* mRNA fold change rarely match those of GFP fluorescence ratios, which is somewhat expected due to the intrinsic differences between the two methodologies. However, we cannot rule out the existence of additional mechanisms affecting *gfp* translation independently of mRNA levels. Indeed, exceptionally some ARCs do not entail changes in *gfp* mRNA levels but decrease significantly GFP fluorescence. Altogether, our results show that polar effects of cassettes are generally observed at the mRNA level.

**Figure 3.**
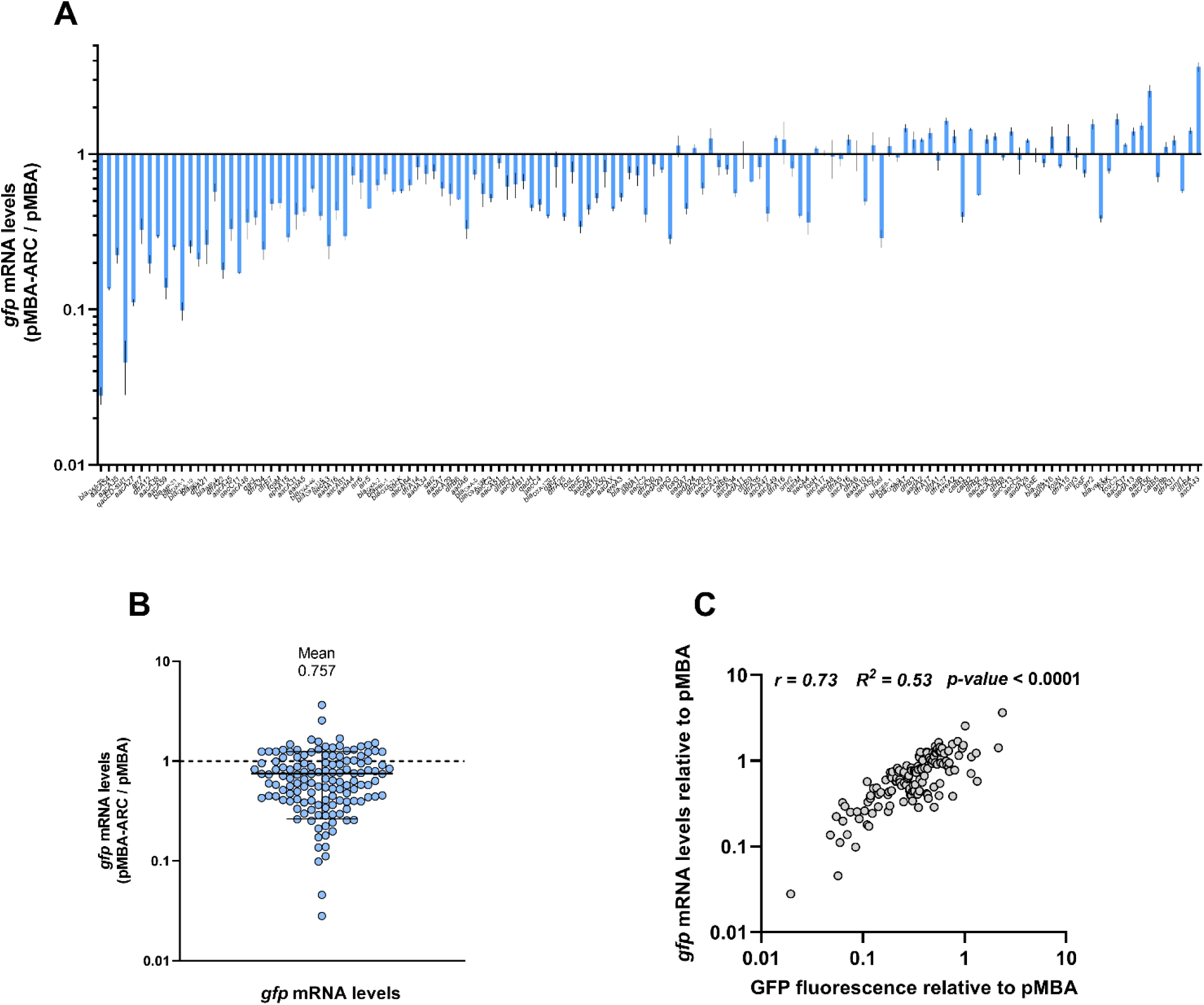
*gfp* mRNA levels of all 136 ARCs. **(A)** *gfp* mRNA levels of all 136 pMBA-ARC strains measured by RT-qPCR normalized to the *gfp* mRNA levels of pMBA control strain (ARCs in the X-axis follow the same order as in Fig. 2A). Bars represent the mean and SD from two to three biological replicates. **(B)** Distribution of the *gfp* mRNA ratios (pMBA-ARC / pMBA) with indication of the mean value and SD. **(C)** Correlation between GFP fluorescence and *gfp* mRNA ratios. *r* indicates the Spearman’s correlation coefficient.

### Negligible role of secondary promoters and *attC* sites in polar effects

The panoply of polar effects observed in this work could be the result of the interplay between several known mechanisms. For instance, cassettes could all have negative polar effects of similar intensity, but some might alleviate them through additional promoters while others intensify them through the presence of transcriptional terminators.

We asked whether additional transcriptional activity could in part explain the high fluorescence levels of a subset of 12 ARCs showing the highest expression levels of the second cassette. These include *qacE*, for which the existence of additional promoters has already been documented ^18,23^; and *ereA2* and *ereA3* that share respectively 94 and 87% identity with *ereA1*, which also contains its own promoter ^14^. We thus deleted the Pc promoter in these 12 variants of pMBA and measured fluorescence to reveal any additional promoters (Fig. 4A). The deletion of the Pc in pMBAø decreased fluorescence 400-fold, to levels similar to a strain without pMBA. Deleting the Pc in this subset confirmed promoter activity in *qacE* and *ereA3*. Other cassettes, like *dfrA1* and *dfrA15* also showed a minor (2-fold) but significantly higher level of fluorescence compared to the pMBAø_ΔPc_ control. Nevertheless, for most cassettes the deletion of the Pc led to a decrease in fluorescence similar to the one observed for pMBAø_ΔPc_. We hence conclude that while additional promoters within certain ARCs might influence the expression of integron arrays, they only account for a limited portion of the overall variation observed in our dataset.

**Figure 4.**
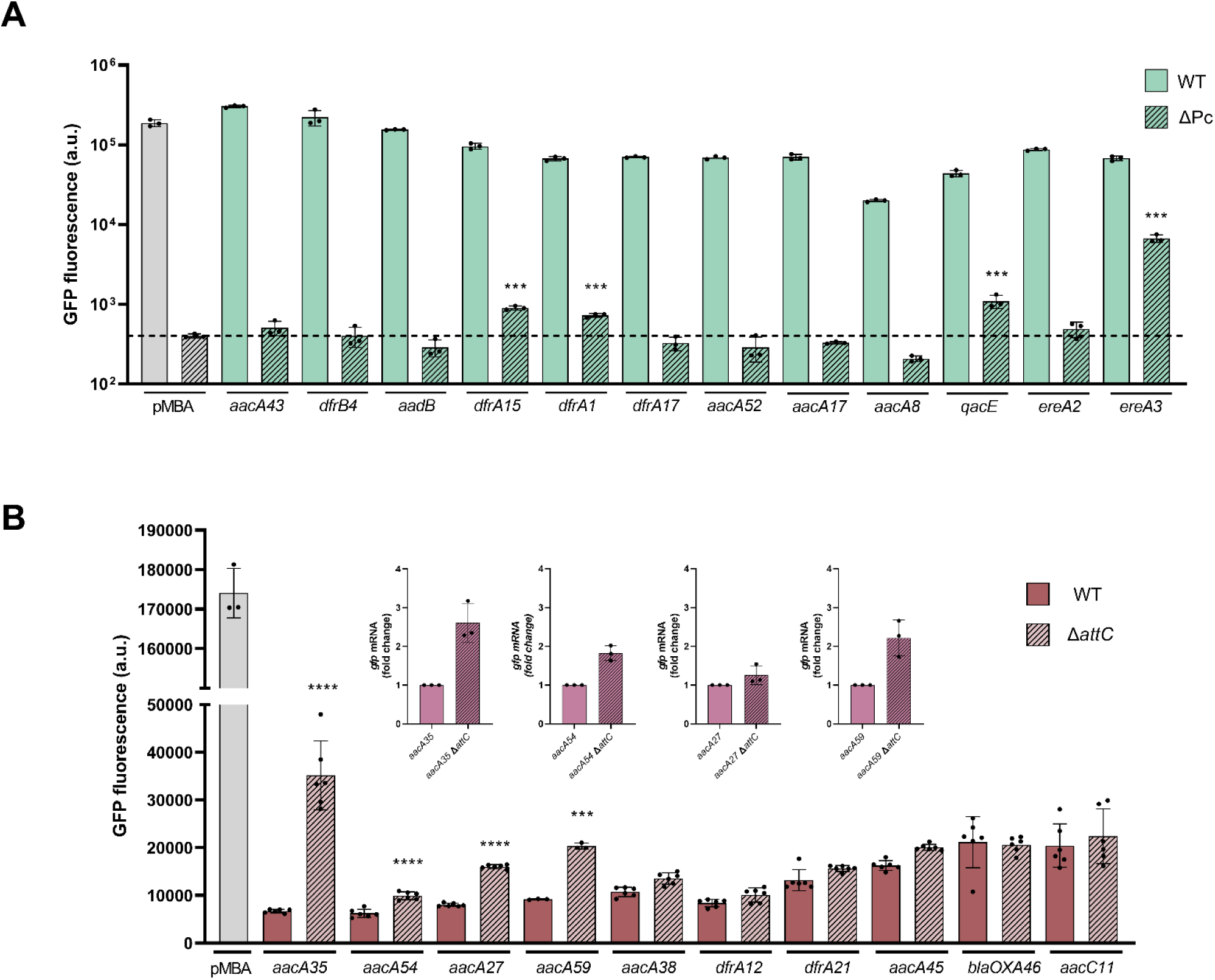
Role of secondary promoters and *attCs* in polar effects. **(A)** Detection of Pc-independent transcriptional activity in a subset of cassettes displaying high GFP expression. Bars depict the GFP fluorescence (arbitrary units) of WT and ΔPc mutants. Dotted line marks the mean value of the pMBAø _ΔPc_ control strain. Statistically significant differences compared to pMBA ΔPc control were determined using unpaired t-test, *** P<0.001. **(B)** Role of *attC* sites in a subset of cassettes displaying low GFP expression. Bars depict GFP fluorescence (arbitrary units) of WT and Δ*attC* mutants. Statistically significant differences between Δ*attC* and WT strains were determined using unpaired t-test, *** P<0.001; **** P<0.0001. Only statistically significant comparisons are shown; (inset) *gfp* mRNA fold change of those Δ*attC* mutants displaying significantly different GFP fluorescence levels.

*attC* recombination sites are sequences located at the 3’ region of each integron cassette. In their recombinogenic form, *attC*s adopt a hairpin structure which is essential for the recognition by the integrase and its recombination. Due to this inherent highly structured conformation, *attC* sites were initially proposed to function as Rho-independent transcriptional terminators ^11^ but later work questioned this view, arguing they rather affect translation of the cassette downstream^13^. We sought to test if *attCs* play a role in the negative polar effects observed, using the subset of 11 ARCs displaying the lowest GFP fluorescence levels. To rule out the influence of plasmid loss in our observations, a phenomenon that could arise from the fitness cost of ARCs, we measured plasmid copy number (PCN) in this subset and pMBAø by qPCR (Supplementary Figure 3). Only pMBA carrying *bla*_OXA-20_ showed a significant reduction in PCN and was thus excluded from further analysis. Deleting *attC* sites in the remaining cassettes reveals no effect on fluorescence for most cases, increasing only mildly the fluorescence in 3 cassettes (Fig. 4B). However, for *aacA35* the deletion of the *attC* results in a large (5-fold) increase in GFP fluorescence. Accordingly, mRNA levels were also higher in 3 out of 4 Δ*attC* strains (Fig. 4B inset). Overall, *attC* removal impacted downstream expression only in a minority of cases and generally in a subtle manner. This suggests that they are not major determinants of polar effects.

### Cassette translation plays a key role on the polar effects in integron arrays

The general absence of promoters within cassettes, together with the lack of repressive effects of *attC* sites suggest that i) these elements only explain a small fraction of polar effects and ii) the underlying mechanism(s) is novel in integrons. We showed above that ARCs significantly affect the mRNA levels of the cassette downstream. Given that the transcription initiation rate originated at the Pc is the same across our collection, we hypothesized that elements within cassettes might either promote transcriptional termination or negatively affect the stability of the polycistronic mRNA ^24^. Considering the critical role of translation in mRNA stability - where translating ribosomes can protect mRNA from degradation ^25–29^ - we examined the translation initiation (TI) rates of each ARC and their correlation with downstream *gfp* mRNA levels (Fig. 5A). We predicted TI rates using the web software RBS Calculator v2.1, which employs a highly accurate thermodynamic model of ribosome-mRNA interactions to estimate the TI rates for a given mRNA sequence ^30^. Our analysis revealed variable TI rates among ARCs and a significant positive correlation between TI rates and *gfp* mRNA levels (ρ = 0.39, *p-value* < 0.0001) (Fig. 5A), highlighting cassette translation as a global determinant in downstream gene expression. Contrarily, we did not observe similar correlations with the codon adaptation index (CAI) of cassettes (ρ = 0.10), or with the overall complexity of secondary structures in their mRNAs (ρ = 0.15), features that have also been shown to be determinants of gene expression and mRNA stability in bacteria ^25,26,31–35^. Interestingly, we observed a negative correlation between the GC content of cassettes and both the TI rates (ρ= -0.37, *p-value* < 0.0001) and *gfp* mRNA levels (ρ= -0.32, *p-value* = 0.001). A low GC content can play a role in gene expression in different ways. When located near the 5’ of ORFs, it can lead to a less structured mRNA, favoring translation initiation ^32,34,36–38^, while along the coding sequence, it may lead to spurious transcription due to AT-rich tracts ^39^. Notably, the correlation between translation rates and the GC content in the 60bp around the start codon was clearly higher than for the whole gene (ρ= -0.49 *vs*. -0.37, *p-value* < 0.0001), suggesting that the impact that GC content has on the polar effects of an ARC is due, at least partly, to its influence on translation initiation rates.

**Figure 5.**
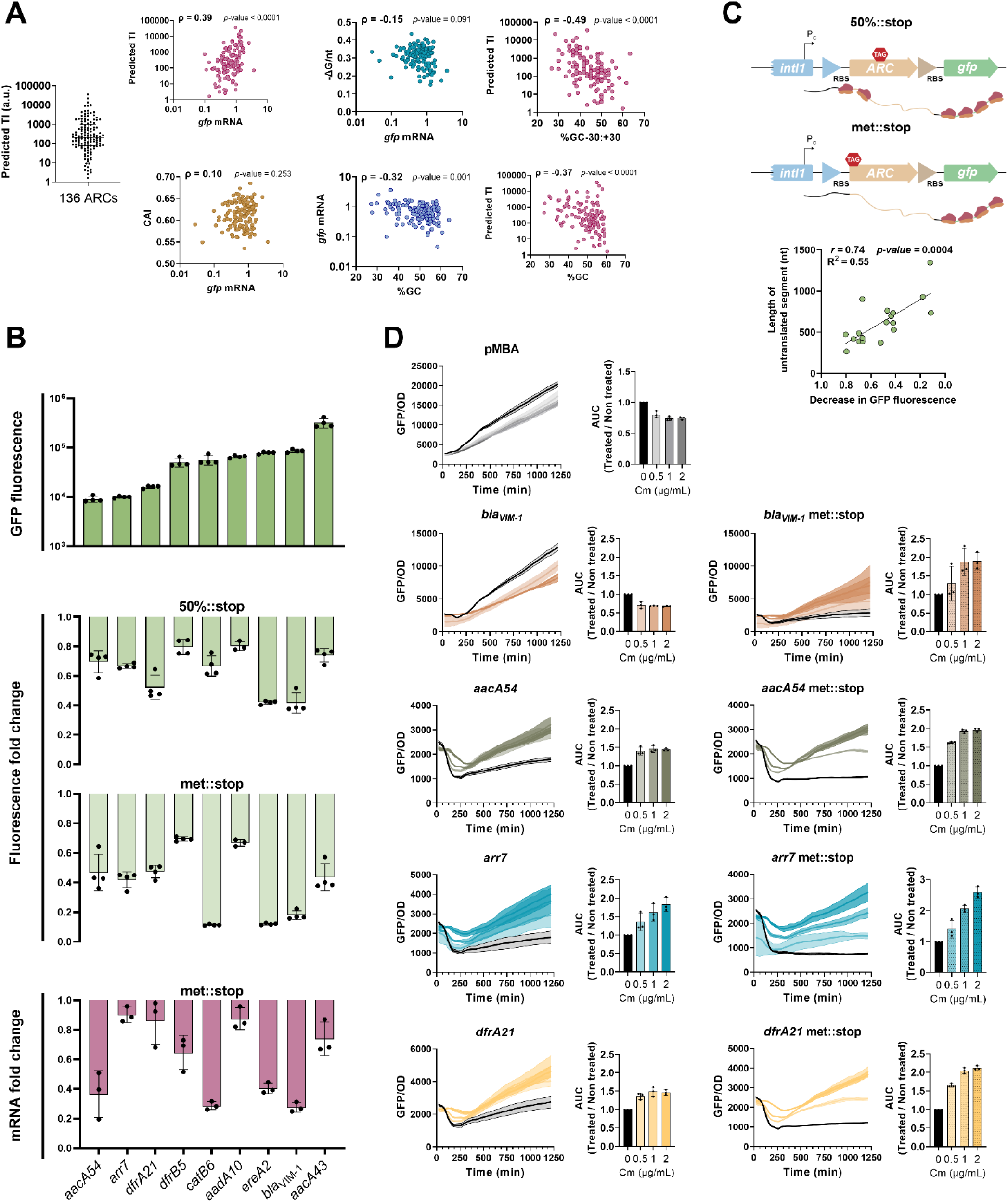
Impact of translation on downstream cassette expression. **(A)** Translation initiation rates of cassettes were predicted using the web tool RBS Calculator 2.1. Correlations between *gfp* mRNA levels of all ARCs and their translation initiation rates (TI); codon adaptation index (CAI) and minimum fold energy of mRNA. Correlations between the GC content of all ARCs and their *gfp* mRNA levels and predicted TI. Spearman’s rank correlation coefficient (**ρ**) and *p-values* are indicated **(B)** GFP fluorescence and *gfp* mRNA levels (fold change) of a subset of ARCs and their respective translation mutants. *met::stop* represents the indicated ARC with the methionine initiation codon replaced by a stop codon. *50%::stop* represents the indicated ARC with a stop codon introduced at the 50% of the CDS. Bars represent the mean and SD from at least three biological replicates. **(C)** Schematic representation of mutants used to study the impact of translation on polar effects in ARCs. Correlation between the length of the untranslated segment and the respective decrease in GFP fluorescence. *r* indicates the Spearman’s correlation coefficient. **(D)** Effect of subinhibitory doses of chloramphenicol on downstream cassette expression. For each indicated strain the left panels display the GFP/OD, with line and shading representing the mean and SD of three biological replicates, respectively. The right panels display the area under the curve (AUC) of the Cm-treated cultures over the non-treated control with bars representing the mean and SD from three biological replicates.

As our data points to the translation of the first cassette as a key driver of the polar effects, we sought to prove it experimentally. We hypothesized that preventing translation initiation of the first cassette, or stopping translation prematurely, should result in a lower expression of the second cassette. To test this, we selected a subset of ARCs from distinct antibiotic families, covering different levels of GFP expression. We hindered translation in these cassettes by i) replacing the ATG initiation codon by a TAG stop codon (met::stop) or ii) by introducing a TAG stop codon in the middle of the CDS (50%::stop). As expected, we observed a general reduction in GFP fluorescence in all untranslated mutants (Fig. 5B). Preventing translation initiation (met::stop mutants) lead to a 2- to 10-fold reduction in downstream cassette expression, while introducing a stop codon in the middle of the CDS (50%::stop) also reduced downstream cassette expression but to a lower extent, suggesting that the sooner translation is prevented, the higher the reduction in downstream expression. Indeed, we find that the length of the untranslated segments in these mutants (i.e., the distance between each stop codon and the ATG of the *gfp* gene), correlates positively with the decrease in fluorescence (Fig. 5C). Notably, in all cases, qRT-PCR of the *gfp* gene shows that downstream cassette mRNA levels also decrease with these mutations, in line with the view that translation interruption affects mRNA levels (Fig. 5B).

To verify that mRNA stability is indeed affected by translation in integrons we sought to prove that stabilization of mRNAs results in an increase in expression of the 2nd cassette. Notably, it has been demonstrated that some translation inhibitors, such as chloramphenicol (Cm), can stabilize bacterial mRNAs ^40–44^. However, because Cm has this dual activity, -inhibiting translation elongation and stabilizing mRNAs-, we hypothesized that the later effect could only be observed in cassettes with low translation levels and low mRNA stability. To test this we measured growth and normalized fluorescence (fluorescence/OD) over time of cultures treated with 0.5, 1 and 2 µg/mL chloramphenicol, (12.5%, 25% and 50% of the MIC, respectively)^20^ (Fig. 5D). The antibiotic effect of Cm was observed on the growth of all strains (Supplementary figure 4) except on the one containing *catB6* which confers high-level resistance to Cm ^20^. As expected, Cm decreased fluorescence in pMBAø, but increased fluorescence of *aacA54*, *arr7* and *dfrA21* cassettes, which are those displaying the lowest fluorescence. In the rest of ARCs, the effect of Cm on fluorescence was intermediate and its sign depended on the initial GFP levels, supporting that there is a balance between the inhibition of translation and the stabilization of unstable mRNAs (Supplementary figure 5 and *bla*_VIM-1_ in Fig. 5D). Additionally, when we prevent translation initiation (met::stop), we observe that Cm treatment results in a higher increase in fluorescence in all cassettes except one (Fig. 5D and Supplementary figure 5). Altogether, these results show that, in general, ARC translation strongly impacts integron array expression by impacting the mRNA levels of downstream cassettes, which are likely influenced by the effect of translation on the stability of the whole mRNA molecule.

### Case-specific analysis of a highly repressive cassette confirms importance of translation

To confirm the importance of translation rates in polar effects, we sought to investigate their role in an extreme case, where repression is maximal and additional mechanisms can be involved. Such is the case of *aacA54,* the most repressive cassette in our collection (it entails 50- and 10-fold decreases of fluorescence and mRNA levels, respectively) (Fig. 2A and 3A). This cassette contains a 25bp 5’ UTR (untranslated region), a 555bp coding sequence, and is one of the very few to contain a long 3’ UTR (162 bp), encompassing the 70bp-long *attC* site (Fig. 6A). We sought to determine the relative importance of translation rates in the polar effects of this cassette, and the presence of other mechanisms.

**Figure 6.**
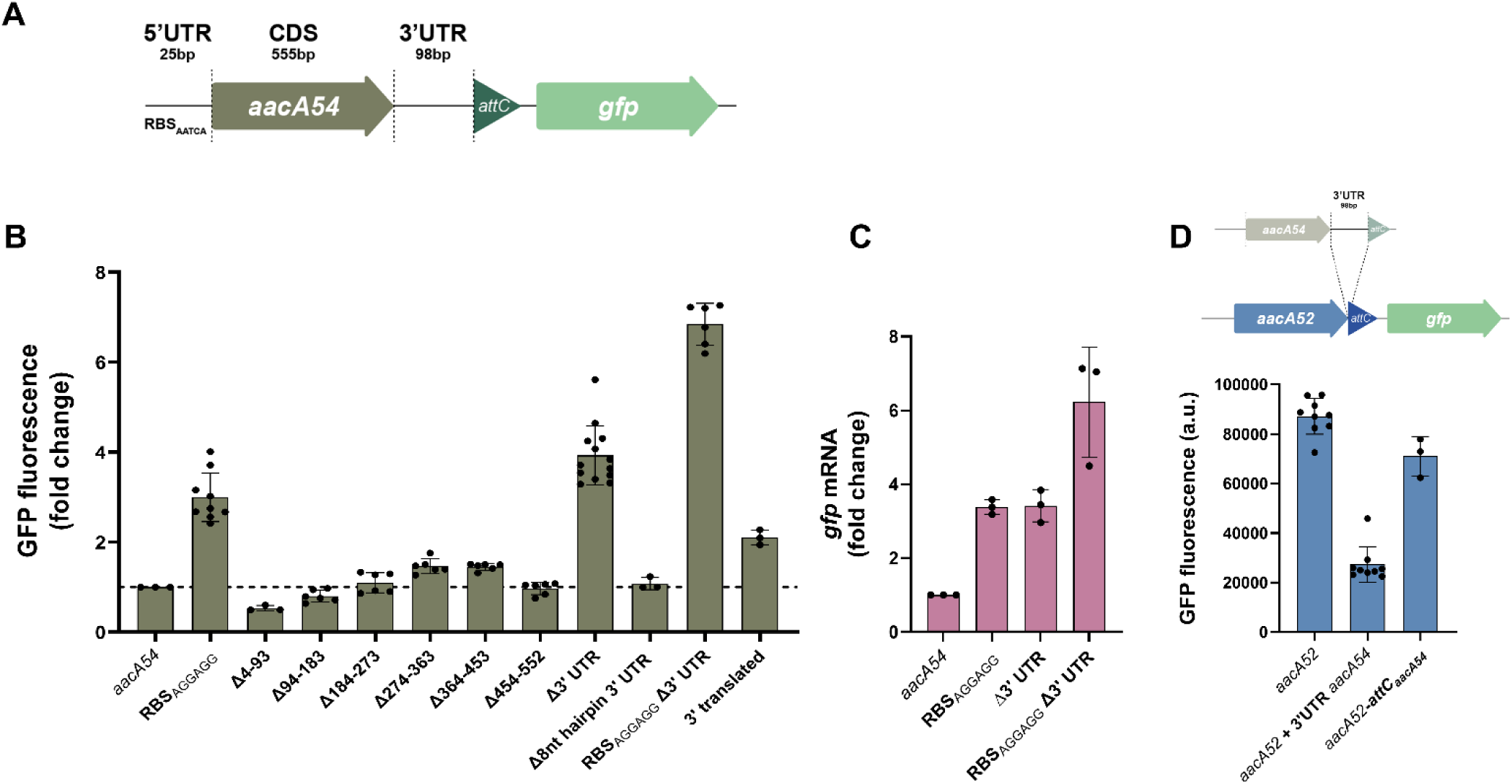
RBS and 3’ UTR of *aacA54* independently govern *gfp* expression. **(A)** Representation of *aacA54* array with each analyzed segment depicted. **(B)** Bar graphics represent the fold change in GFP fluorescence of each modified strain relative to the original *aacA54* strain. **(C)** *gfp* mRNA fold change of the indicated *aacA54* mutants. Bars represent the mean and SD from at least three independent biological replicates. **(D)** Representation of *aacA52* modified strain now including the 98bp segment from *aacA54*. Bar graphics represent GFP fluorescence levels (arbitrary units) of original or modified *aacA52* strains.

The 5’ UTR region of genes contains the Ribosome Binding Site (RBS) and other signals (like mRNA structures or A-rich tracts) that collectively control their translation initiation rate ^37,45–47^. In this sense, *aacA54* harbors a Shine-Dalgarno (SD) motif, AATCAA, that is far from the canonical AGGAGG. To assess if this weak RBS might result in a low translation rate for *aacA54* and a low stability of mRNA, we modified the SD to the canonical AGGAGG. Notably, the resistance levels against four different aminoglycosides increased 2- to 4-fold, confirming an increased translation rate in this mutant (Supplementary figure 6). This change in the RBS of the first cassette increased 3-fold the expression (Fig. 6B) and the mRNA levels of the second cassette (the *gfp* gene) (Fig. 6C) which confirms the important role of translation rates.

Additionally, we searched for elements within the CDS of *aacA54* that could lower the expression of the second cassette, such as stretches of rare codons, known to slow down translation ^25,32^. By performing serial 90bp deletions covering the whole length of *aacA54*, preserving the start and stop codons as well as the reading frame, we found no changes in fluorescence (Fig. 6B), indicating that the CDS does not affect downstream expression.

As mentioned before, *aacA54* contains a 98bp long 3’ UTR that separates the stop codon from the *attC* site, contrarily to the majority of the other ARCs where the *attC* overlaps with the stop codon of the CDS. We hypothesized that the presence of this region could negatively affect the expression levels of the next cassette, either due to the presence of transcriptional terminators^48^ or an untranslated stretch of mRNA. Indeed, the deletion of the UTR resulted in a 4-fold increase in fluorescence and a similar increase in *gfp* mRNA levels (Fig. 6B and 6C). When combined with a canonical RBS, the effect was additive both at the fluorescence and the mRNA levels (Fig. 6B and 6C), suggesting that both elements act independently. Moreover, when we introduced this 3’ UTR between the stop codon and the *attC* site of *aacA52*, a closely related cassette with high initial fluorescence levels, we observed a 4-fold reduction in fluorescence (Fig. 6D). Instead, replacing only the *attC_aacA52_* site with *attC_aacA54_* caused a minor reduction in fluorescence. We identified a small hairpin-like structure that could be acting as an intrinsic transcriptional terminator (Supplementary figure 7). However, removing 8 nucleotides essential for this structure did not affect fluorescence levels, ruling out the role of this sequence as a terminator (Fig. 6B). We then mutated the canonical stop codon in *aacA54* so that translation is maintained in frame throughout all the 3’ region until the *attC*, where it terminates at a new stop codon. This led to a 2-fold increase in fluorescence, which agrees, in part, with the view that lack of translation at the 3’ UTR lowers downstream expression (Fig. 6B).

In summary, translation rates are a key element in polar effects even in extremely repressive cassettes.

## DISCUSSION

The current working model of integrons puts forward that cassettes in an array follow a passive gradient of expression that depends on their distance to the Pc. Hence, for any given integron platform (i.e. assuming the same Pc variant), the effect of cassette position is considered as the main determinant of cassette expression. However, the potential impact of ARCs on the expression of cassettes downstream, and consequently the resistance levels of integrons, has been largely overlooked in the field. In this work, we have used the pMBA collection^20^ to show that the expression of a cassette is strongly determined by the identity of the cassette that precedes it, to the point of masking the “distance-to-Pc” effect.

Our results show that translation of cassettes in first position is key in the expression of second cassettes. While we clearly show this with experimental data using cassettes from different antibiotic families, the translation initiation rates predicted bioinformatically for all ARCs correlate only mildly with their polar effects. We suspect that other factors not accounted for in these predictions may also be affecting translation rates for certain cassettes. For example, some cassettes in our collection are predicted to have a very low translation initiation rate, yet this does not necessarily mean they are not translated. Indeed, it has been suggested that cassettes without a proper translation initiation region may be translated due to lateral diffusion of ribosomes from translating ORF-11 or ORF-17, which are small ORFs encoded at the *attI* site ^49,50^. It is also possible that some cassettes in our collection contain misannotated initiation codons that bias translation rates.

We propose that the link between ARC’s translation and polar effects lies in the intimate association between translation and mRNA stability. Indeed, translation may modulate the overall stability of the transcript generated at the Pc, by protecting it against degradation by RNases. Thus, considering our results showing that different ARCs have different translation rates, one can also assume they are likely to display different transcript stabilities. The fact that chloramphenicol, as an RNA stabilizing agent, increased expression downstream of poorly translated cassettes further supports this view.

Nevertheless, translation is likely not the only factor producing polar effects in integron arrays. Indeed, we have found some exceptions in our findings that support the occurrence of other phenomena, such as cases of known and novel promoter activity in *ereA3*, *dfrA15*, *dfrA1* and *qacE*; and the negative effect of the *attC* site of *aacA35*. As for the mechanism through which some *attC*s lower downstream expression, one can hypothesize that the highly structured form of *attC*s may be facilitated by the lack of ribosome trafficking in cassettes with low translation rates, which could potentially leave the 5’ region of the *gfp “*cassette” temporarily more exposed to ribonucleolytic attack.

Our finding that the identity of cassettes ultimately controls polar effects in integrons may have important consequences in the clinical context. A limitation of our study is that we used a multicopy vector, which likely results in higher absolute MIC values than those in natural environments, as discussed in ^20^. Nonetheless, we believe the described polar effects should exist independently of copy number or Pc variant in natural integrons. The fact that a given ARC may confer different levels of resistance, depending on the preceding cassette has important consequences in co-selection phenomena. Indeed, antibiotic combination therapies may select for arrays whose cassette order allows for the expression of the entire array. Moreover, considering the fitness costs entailed by the majority of antibiotic resistance genes, and how it relates to the expression levels of the AR gene ^51,52^, cassette order can optimize the cost of the array. Interestingly, some resistance genes -such as trimethoprim resistance *dfrs*-have an almost digital phenotype, conferring high resistance even at very low expression levels. Hence, one could imagine a situation in which a cassette represses the expression of the next ARC in the array, decreasing its cost, but maintaining its resistance phenotype. In other words, in the light of our findings, integrons can optimize the tradeoff between fitness cost and function of antibiotic resistance genes.

This work changes the paradigm of expression in integrons, and -together with other works showing the presence of promoters in cassettes-puts forward a more complex scenario. In this new model, inferring the levels of expression of a given cassette is extremely challenging and needs a case-by-case assessment, especially if preceded by cassettes not found in our collection.

## MATERIAL AND METHODS

### Bacterial strains, plasmids, and culture conditions

*Escherichia coli* MG1655 strains and plasmids used in this study are based on the previously published pMBA collection ^20^ and are listed in Supplementary Table 1. All strains were cultured in liquid Müeller Hinton (MH; Oxoid, UK) media, at 37°C with agitation at 200 rpm or solid lysogeny broth (LB) agar (1.5%) (BD, France). Zeocin (Zeo) (Invitrogen, USA) was added at 100µg/mL for plasmid maintenance. All pMBA collection-derived mutants were obtained by backbone amplification using primers designed to achieve the desire modifications followed by Gibson assembly ^53^ (Supplementary Table 1).

### Flow cytometry analysis of fluorescence

Strains containing pMBA ARC-gfp plasmids were previously streaked on LB-solid medium with 100µg/mL Zeo. Three independent colonies were then inoculated in 200µL MH liquid medium supplemented with 100µg/mL Zeocin and incubated at 37°C with agitation for 20h. The next day, cultures were diluted 1:100 in 200µL MH+Zeo and incubated for 2h to reach exponential phase. At this point, cultures were diluted 1:20 in filtered saline solution (NaCl 0.9%) and fluorescent intensity was measured by flow cytometry using a CytoFLEX-S cytometer (Beckman Coulter, USA). The measurement of each biological replicate is the result of the mean of the fluorescence intensity of 30 000 events per sample. Data processing was performed with Cytexpert software.

### Quantitative Reverse Transcription PCR of *gfp* gene

For RNA extraction, overnight cultures of three biological replicates of each strain were diluted 1:100 in MH medium supplemented with zeocin (100 µg/mL) and grown in 96 well plates with agitation at 37°C for approximately 2 – 3 hours, until reaching exponential phase. Total RNA was extracted using a KingFisher Flex automated system with the MagMAX mirVana Total RNA Isolation Kit from Applied Biosystems, according to manufacturer’s instructions. The purity and concentration of total RNA was determined by spectrophotometry (BioSpectrometer). For cDNA synthesis, 200ng of RNA was used for reverse transcription using the QuantiTect Reverse Transcription Kit (Qiagen) according to manufacturer’s instructions. A negative control without Reverse Transcriptase was included to exclude possible gDNA carryover. cDNA was diluted 1:100 and 1µL of this dilution was used for quantitative PCR (qPCR). qPCR was performed in QuantStudio 3 Real-Time PCR system (Applied Biosystems) with the QuantiTect Multiplex PCR kit (Qiagen) according to manufacturer’s instructions. Primers and Probes used in the mix are listed in SUP TABLE. PCR thermocycling conditions contained an initial stage of 2 min at 50°C and 15 min at 95°C followed by 42 cycles of amplification (1 min at 94°C and 1 min at 60°C). Negative controls included both a reaction containing water instead of template and a reverse transcriptase-free reaction. The relative abundance of *gfp* transcripts was normalized to that of the housekeeping gene *rssA*^54^ using the 2^-ΔΔCt^ method ^55^.

### Plasmid copy number (PCN) assessment

PCN was assessed as in ^56^. Briefly, overnight cultures of three biological replicates of each strain were diluted 1:100 in MH medium supplemented with zeocin (100 µg/mL) and grown in 96 well plates with agitation at 37°C for approximately 2 hours, until reaching exponential phase. 50µL were then collected and briefly centrifuged for 2 minutes. The supernatant was then discarded, the pellet resuspended in molecular biology grade water and then boiled at 98°C for 10 minutes. After a brief centrifugation, 30µL were collected and then diluted 1:10. 1 µL of this dilution was then used as template for the exact same qPCR reaction described above.

### Growth curves and GFP measurement of chloramphenicol-treated cultures

Three independent colonies of each strain were inoculated in MH+Zeo and incubated at 37°C with agitation for 20h. Cultures were then diluted 1:1000 in fresh MH+Zeo media containing or not subinhibitory concentrations of chloramphenicol (0.5, 1 and 2µg/mL). Growth (OD_600_) and GFP fluorescence (measured at 488nm wavelength) were followed for 20h using a Biotek Synergy HTX plate reader. Measures were taken every 20 minutes with prior shaking.

### Determination of minimum inhibitory concentrations (MIC)

The MIC of ertapenem, cefaclor or chloramphenicol of strains containing *bla_VIM-1_* or *catB6* cassettes were determined as in ^20^. Briefly, 10^5^ colony forming units (CFUs) were inoculated in 200μL of fresh MH with doubling dilutions of each selected antibiotic in 96-well plates and incubated overnight at 37°C in static conditions. After 24 hours plates were analyzed and MIC values were established as the lowest concentration in which visible growth could not be observed.

## ACKNOWLEDGEMENTS

We would like to thank José R. Penadés for helpful discussions and for the critical reading of the manuscript. We are grateful to all the members of the MBA lab for helpful discussion.

## AUTHOR CONTRIBUTIONS

AC: conceptualization, methodology, formal analysis, validation, and writing-original draft. AH: conceptualization, methodology, formal analysis, validation. FTR: methodology, formal analysis, validation. LGP, methodology, formal analysis, validation; EV, methodology, formal analysis. AB, methodology; TGS methodology. JAE conceptualization, formal analysis, validation, and writing the final manuscript. All authors read amended and approved the final version of the manuscript.

## SUPPLEMENTARY INFORMATION

**Table S1**: Bacterial strains, Plasmids and Primers used in this study.

**Supplementary figure 1.**
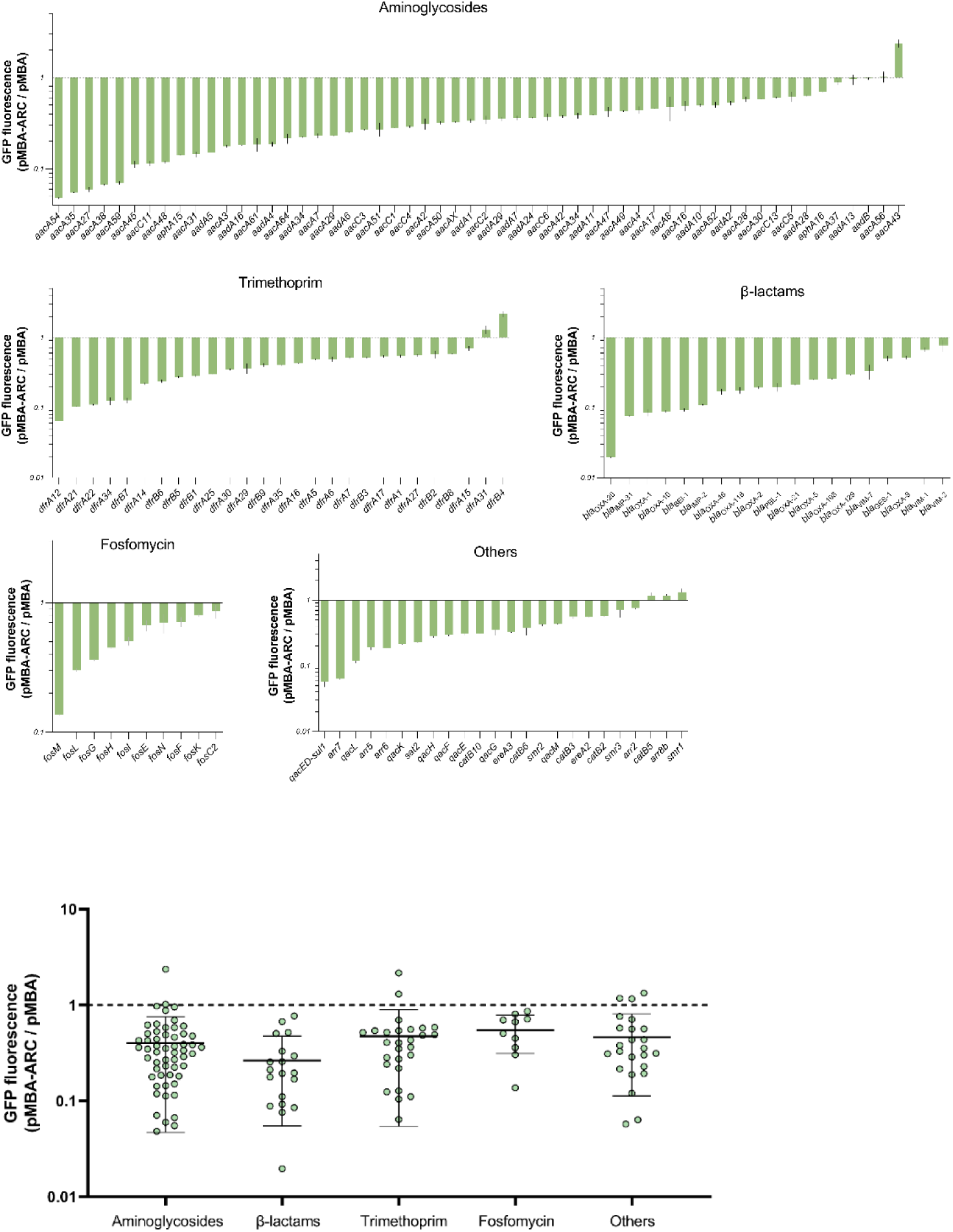
Polar effects by antibiotic families. GFP fluorescence of all 136 pMBA-ARC strains measured by flow cytometry normalized to fluorescence of pMBA control strain. Bars represent the mean and SD from three independent biological replicates. Distribution of the GFP fluorescence ratios (pMBA-ARC / pMBA) with indication of the mean value and SD.

**Supplementary figure 2.**
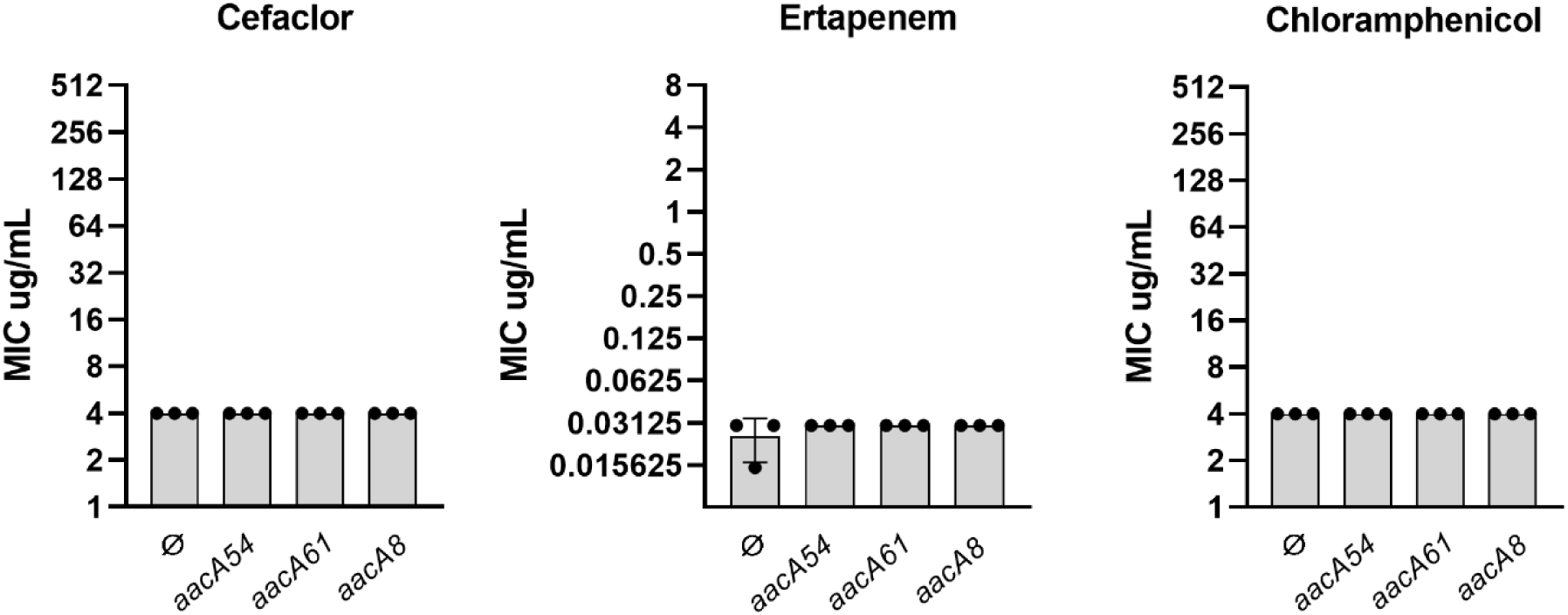
Resistance levels conferred by pMBAø or pMBA carrying *aacA54*, *aacA61* or *aacA8* to cefaclor, ertapenem and chloramphenicol. Bars depict the mean and SD of the MIC values of three independent biological replicates.

**Supplementary figure 3.**
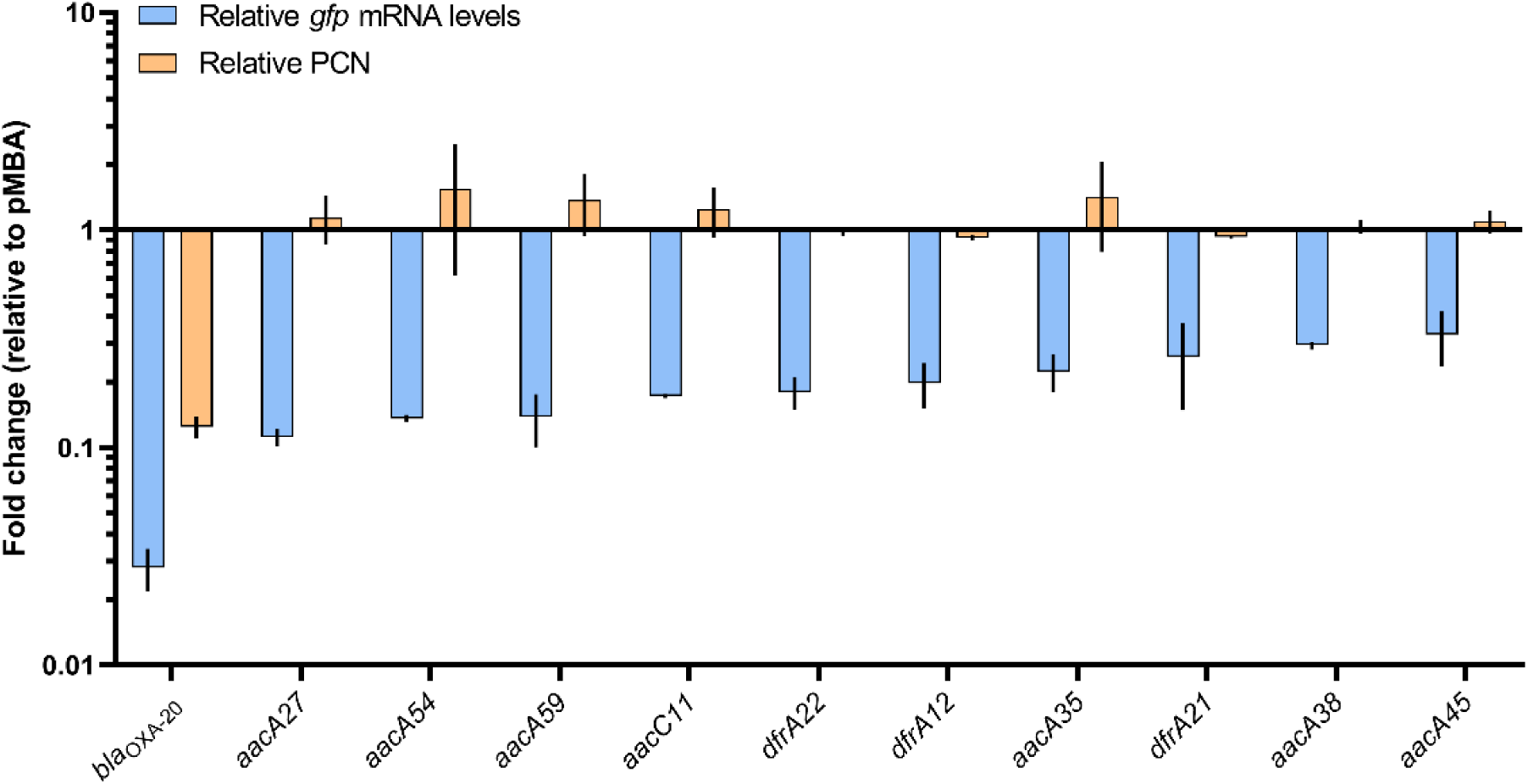
Relative plasmid copy number (PCN) and relative *gfp* mRNA levels of low *gfp* strains. PCN was determined by qPCR using total lysate DNA (see Material and Methods). Bars represent the mean and SD from two to three independent biological replicates.

**Supplementary figure 4.**
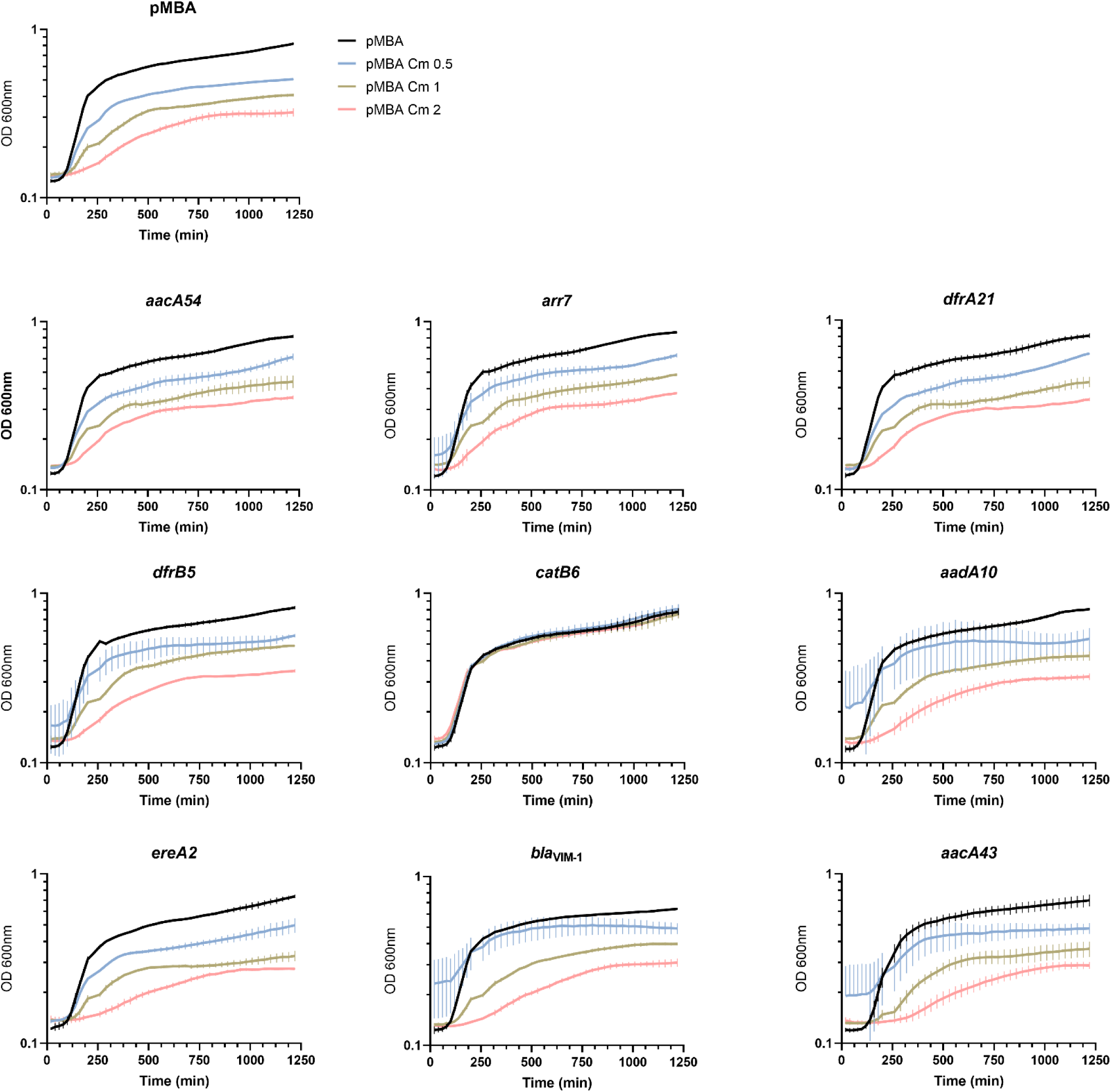
Growth (OD_600_) of the indicated strains in absence or presence of 0.5, 1, and 2 µg/mL chloramphenicol. Lines indicate the mean value and error bars represent the SD of three biological replicates.

**Supplementary figure 5.**
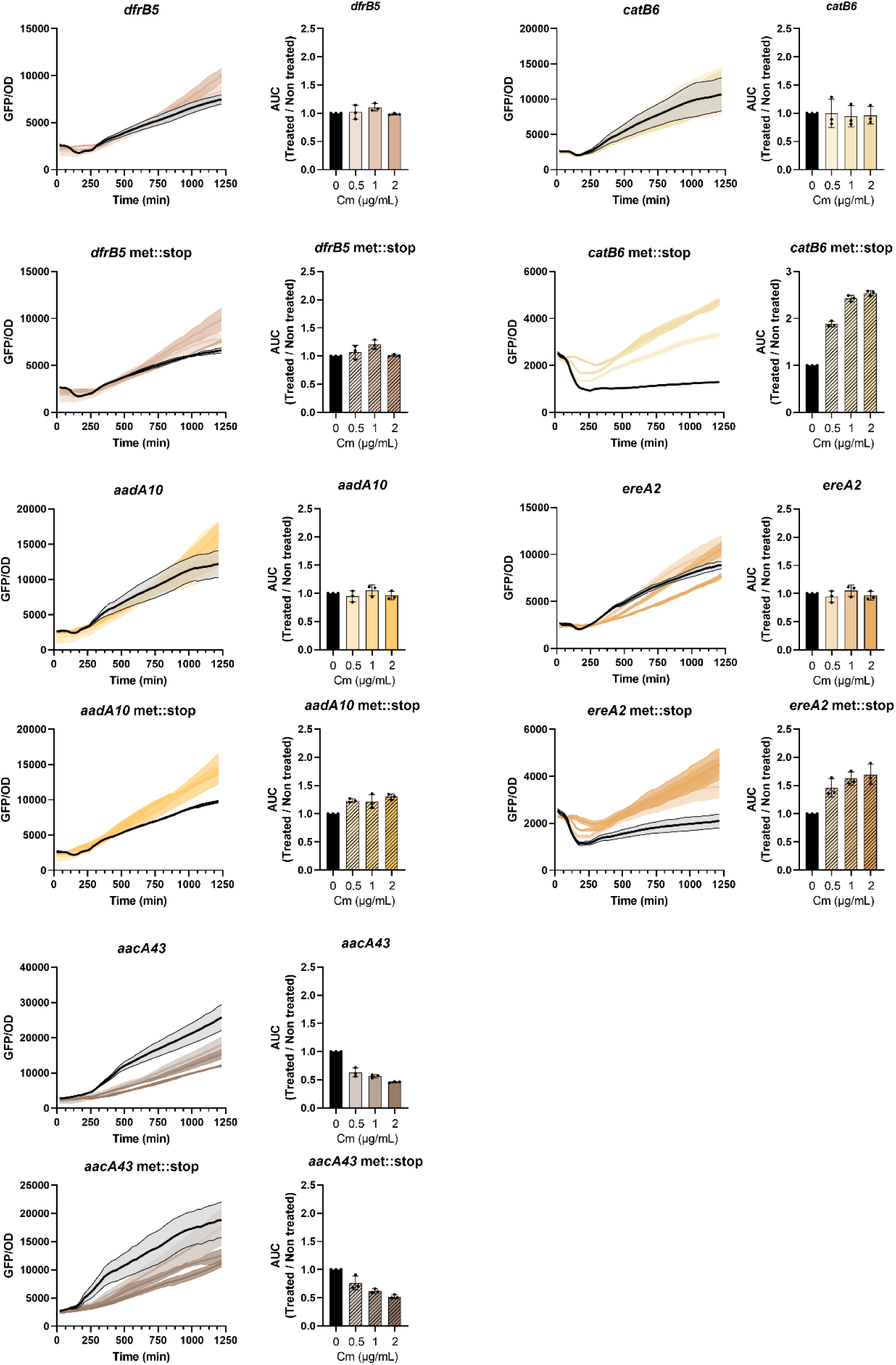
Effect of subinhibitory doses of chloramphenicol on downstream cassette expression. For each indicated strain the left panels display the GFP/OD, with line and shading representing the mean and SD of three biological replicates, respectively. The right panels display the area under the curve (AUC) of the Cm-treated cultures over the non-treated control with bars representing the mean and SD from three independent biological replicates.

**Supplementary figure 6.**
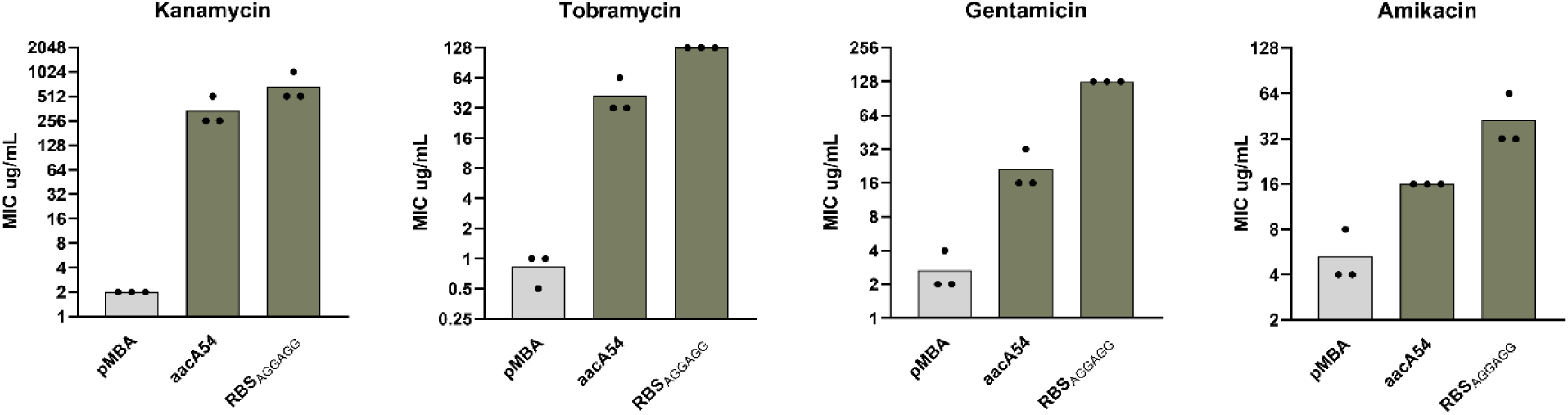
Resistance levels conferred by pMBAø or pMBA carrying the indicated arrays to the four aminoglycosides. Bars depict the mean and SD of the MIC values of three independent biological replicates.

**Supplementary figure 7.**
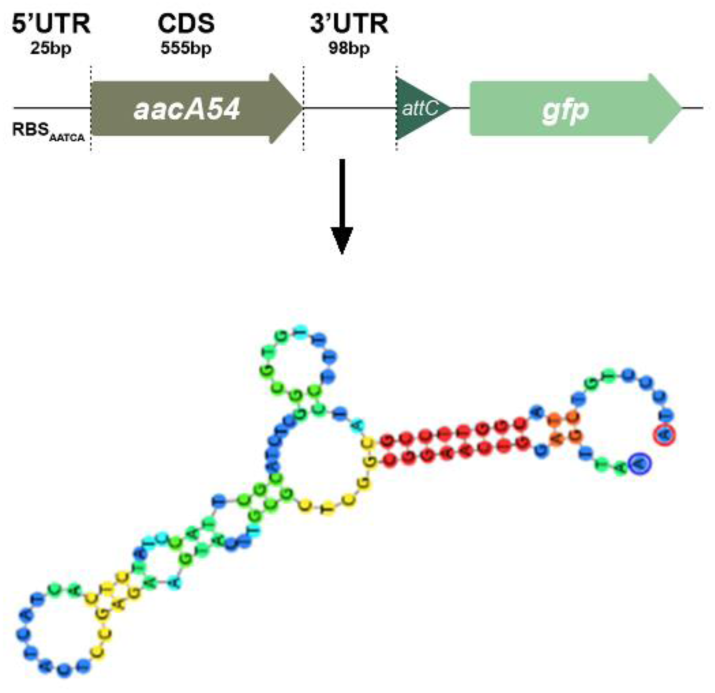
Representation of the putative hairpin-like structure of 3’ UTR in *aacA54*.

## REFERENCES

1. Murray, C. J. L. et al. Global burden of bacterial antimicrobial resistance in 2019: a systematic analysis. The Lancet 399, 629–655 (2022).

2. Gillings, M. et al. The evolution of class 1 integrons and the rise of antibiotic resistance. J. Bacteriol. 190, 5095–5100 (2008).

3. Gillings, M. R. Class 1 integrons as invasive species. Curr. Opin. Microbiol. 38, 10–15 (2017).

4. Partridge, S. R., Tsafnat, G., Coiera, E. & Iredell, J. R. Gene cassettes and cassette arrays in mobile resistance integrons. FEMS Microbiol. Rev. 33, 757–784 (2009).

5. Moura, A. et al. INTEGRALL: a database and search engine for integrons, integrases and gene cassettes. Bioinformatics 25, 1096–1098 (2009).

6. Guerin, É. et al. The SOS Response Controls Integron Recombination. Science 324, 1034–1034 (2009).

7. Barraud, O. & Ploy, M.-C. Diversity of Class 1 Integron Gene Cassette Rearrangements Selected under Antibiotic Pressure. J. Bacteriol. 197, 2171–2178 (2015).

8. Escudero, J. A., Loot, C., Nivina, A. & Mazel, D. The Integron: Adaptation On Demand. Microbiol. Spectr. 3, 10.1128/microbiolspec.mdna3-0019–2014 (2015).

9. Lacotte, Y., Ploy, M.-C. & Raherison, S. Class 1 integrons are low-cost structures in Escherichia coli. ISME J. 11, 1535–1544 (2017).

10. Souque, C., Escudero, J. A. & MacLean, R. C. Integron activity accelerates the evolution of antibiotic resistance. eLife 10, e62474 (2021).

11. Collis, C. M. & Hall, R. M. Expression of antibiotic resistance genes in the integrated cassettes of integrons. Antimicrob. Agents Chemother. 39, 155–162 (1995).

12. da Fonseca, E. L., Freitas, F. d. S. & Vicente, A. C. P. Pc promoter from class 2 integrons and the cassette transcription pattern it evokes. J. Antimicrob. Chemother. 66, 797–801 (2011).

13. Jacquier, H., Zaoui, C., Sanson-le Pors, M.-J., Mazel, D. & Berçot, B. Translation regulation of integrons gene cassette expression by the *attC* sites. Mol. Microbiol. 72, 1475–1486 (2009).

14. Biskri, L. & Mazel, D. Erythromycin esterase gene ere(A) is located in a functional gene cassette in an unusual class 2 integron. Antimicrob. Agents Chemother. 47, 3326–3331 (2003).

15. Bissonnette, L., Champetier, S., Buisson, J. P. & Roy, P. H. Characterization of the nonenzymatic chloramphenicol resistance (cmlA) gene of the In4 integron of Tn1696: similarity of the product to transmembrane transport proteins. J. Bacteriol. 173, 4493 (1991).

16. da Fonseca, É. L. & Vicente, A. C. P. Functional Characterization of a Cassette-Specific Promoter in the Class 1 Integron-Associated qnrVC1 Gene. Antimicrob. Agents Chemother. 56, 3392–3394 (2012).

17. Guérout, A.-M. et al. Characterization of the phd-doc and ccd Toxin-Antitoxin Cassettes from Vibrio Superintegrons. J. Bacteriol. 195, 2270–2283 (2013).

18. Poirel, L. et al. Characterization of Class 1 Integrons from Pseudomonas aeruginosa That Contain the blaVIM-2Carbapenem-Hydrolyzing β-Lactamase Gene and of Two Novel Aminoglycoside Resistance Gene Cassettes. Antimicrob. Agents Chemother. 45, 546–552 (2001).

19. Blanco, P. et al. Identification of promoter activity in gene-less cassettes from Vibrionaceae superintegrons. Nucleic Acids Res. gkad1252 (2024) doi:10.1093/nar/gkad1252.

20. Hipólito, A., García-Pastor, L., Vergara, E., Jové, T. & Escudero, J. A. Profile and resistance levels of 136 integron resistance genes. Npj Antimicrob. Resist. 1, 1–12 (2023).

21. Hipólito, A. et al. The expression of aminoglycoside resistance genes in integron cassettes is not controlled by riboswitches. Nucleic Acids Res. 50, 8566–8579 (2022).

22. Starikova, I. et al. A Trade-off between the Fitness Cost of Functional Integrases and Long-term Stability of Integrons. PLoS Pathog. 8, e1003043 (2012).

23. Naas, T., Mikami, Y., Imai, T., Poirel, L. & Nordmann, P. Characterization of In53, a Class 1 Plasmid- and Composite Transposon-Located Integron of Escherichia coli Which Carries an Unusual Array of Gene Cassettes. J. Bacteriol. 183, 235–249 (2001).

24. Güell, M., Yus, E., Lluch-Senar, M. & Serrano, L. Bacterial transcriptomics: what is beyond the RNA horiz-ome? Nat. Rev. Microbiol. 9, 658–669 (2011).

25. Boël, G. et al. Codon influence on protein expression in E. coli correlates with mRNA levels. Nature 529, 358–363 (2016).

26. Dar, D. & Sorek, R. Extensive reshaping of bacterial operons by programmed mRNA decay.PLOS Genet. 14, e1007354 (2018).

27. Deana, A. & Belasco, J. G. Lost in translation: the influence of ribosomes on bacterial mRNA decay. Genes Dev. 19, 2526–2533 (2005).

28. Duviau, M.-P. et al. When translation elongation is impaired, the mRNA is uniformly destabilized by the RNA degradosome, while the concentration of mRNA is altered along the molecule. Nucleic Acids Res. 51, 2877–2890 (2023).

29. Viegas, S. C., Apura, P., Martínez-García, E., de Lorenzo, V. & Arraiano, C. M. Modulating Heterologous Gene Expression with Portable mRNA-Stabilizing 5′-UTR Sequences. ACS Synth. Biol. 7, 2177–2188 (2018).

30. Reis, A. C. & Salis, H. M. An Automated Model Test System for Systematic Development and Improvement of Gene Expression Models. ACS Synth. Biol. 9, 3145–3156 (2020).

31. Burkhardt, D. H. et al. Operon mRNAs are organized into ORF-centric structures that predict translation efficiency. eLife 6, e22037 (2017).

32. Cambray, G., Guimaraes, J. C. & Arkin, A. P. Evaluation of 244,000 synthetic sequences reveals design principles to optimize translation in Escherichia coli. Nat. Biotechnol. 36, 1005–1015 (2018).

33. Cetnar, D. P. & Salis, H. M. Systematic Quantification of Sequence and Structural Determinants Controlling mRNA stability in Bacterial Operons. ACS Synth. Biol. 10, 318–332 (2021).

34. Gu, W., Zhou, T. & Wilke, C. O. A Universal Trend of Reduced mRNA Stability near the Translation-Initiation Site in Prokaryotes and Eukaryotes. PLOS Comput. Biol. 6, e1000664 (2010).

35. Kudla, G., Murray, A. W., Tollervey, D. & Plotkin, J. B. Coding-Sequence Determinants of Gene Expression in Escherichia coli. Science 324, 255–258 (2009).

36. Allert, M., Cox, J. C. & Hellinga, H. W. Multifactorial determinants of protein expression in prokaryotic open reading frames. J. Mol. Biol. 402, 905–918 (2010).

37. Komarova, A. V., Tchufistova, L. S., Dreyfus, M. & Boni, I. V. AU-Rich Sequences within 5′ Untranslated Leaders Enhance Translation and Stabilize mRNA in Escherichia coli. J. Bacteriol. 187, 1344–1349 (2005).

38. Lenz, G., Doron-Faigenboim, A., Ron, E. Z., Tuller, T. & Gophna, U. Sequence Features of E. coli mRNAs Affect Their Degradation. PLOS ONE 6, e28544 (2011).

39. Warman, E. A., Singh, S. S., Gubieda, A. G. & Grainger, D. C. A non-canonical promoter element drives spurious transcription of horizontally acquired bacterial genes. Nucleic Acids Res. 48, 4891–4901 (2020).

40. Lopez, P. J., Marchand, I., Yarchuk, O. & Dreyfus, M. Translation inhibitors stabilize Escherichia coli mRNAs independently of ribosome protection. Proc. Natl. Acad. Sci. 95, 6067–6072 (1998).

41. Lundberg, U., Nilsson, G. & von Gabain, A. The differential stability of the Escherichia coli ompA and bla mRNA at various growth rates is not correlated to the efficiency of translation. Gene 72, 141–149 (1988).

42. Pato, M. L., Bennett, P. M. & Von Meyenburg, K. Messenger Ribonucleic Acid Synthesis and Degradation in Escherichia coli During Inhibition of Translation. J. Bacteriol. 116, 710–718 (1973).

43. Richards, J., Luciano, D. J. & Belasco, J. G. Influence of translation on RppH-dependent mRNA degradation in Escherichia coli. Mol. Microbiol. 86, 1063–1072 (2012).

44. Vargas-Blanco, D. A. & Shell, S. S. Regulation of mRNA Stability During Bacterial Stress Responses. Front. Microbiol. 11, (2020).

45. Chen, F., Cocaign-Bousquet, M., Girbal, L. & Nouaille, S. 5’UTR sequences influence protein levels in Escherichia coli by regulating translation initiation and mRNA stability. Front. Microbiol. 13, (2022).

46. Duan, Y. et al. Deciphering the Rules of Ribosome Binding Site Differentiation in Context Dependence. ACS Synth. Biol. 11, 2726–2740 (2022).

47. Ma, J., Campbell, A. & Karlin, S. Correlations between Shine-Dalgarno Sequences and Gene Features Such as Predicted Expression Levels and Operon Structures. J. Bacteriol. 184, 5733– 5745 (2002).

48. Menendez-Gil, P. & Toledo-Arana, A. Bacterial 3′UTRs: A Useful Resource in Post-transcriptional Regulation. Front. Mol. Biosci. 7, 617633 (2021).

49. Hanau-Berçot, B., Podglajen, I., Casin, I. & Collatz, E. An intrinsic control element for translational initiation in class 1 integrons. Mol. Microbiol. 44, 119–130 (2002).

50. Papagiannitsis, C. C., Tzouvelekis, L. S., Tzelepi, E. & Miriagou, V. attI1-Located Small Open Reading Frames ORF-17 and ORF-11 in a Class 1 Integron Affect Expression of a Gene Cassette Possessing a Canonical Shine-Dalgarno Sequence. Antimicrob. Agents Chemother. 61, 10.1128/aac.02070-16 (2017).

51. Vogwill, T. & MacLean, R. C. The genetic basis of the fitness costs of antimicrobial resistance: a meta-analysis approach. Evol. Appl. 8, 284–295 (2015).

52. Rajer, F. & Sandegren, L. The Role of Antibiotic Resistance Genes in the Fitness Cost of Multiresistance Plasmids. mBio 13, e03552–21 (2022).

53. Gibson, D. G. et al. Enzymatic assembly of DNA molecules up to several hundred kilobases. Nat. Methods 6, 343–345 (2009).

54. Peng, S., Stephan, R., Hummerjohann, J. & Tasara, T. Evaluation of three reference genes of Escherichia coli for mRNA expression level normalization in view of salt and organic acid stress exposure in food. FEMS Microbiol. Lett. 355, 78–82 (2014).

55. Livak, K. J. & Schmittgen, T. D. Analysis of Relative Gene Expression Data Using Real-Time Quantitative PCR and the 2−ΔΔCT Method. Methods 25, 402–408 (2001).

56. Rodriguez-Beltran, J. et al. Multicopy plasmids allow bacteria to escape from fitness trade-offs during evolutionary innovation. *Nat*. Ecol. Evol. 2, 873 (2018).

